# Uncovering genetic determinants of rice root anatomy using genome wide analysis

**DOI:** 10.64898/2026.10.13.758744

**Authors:** Hien Linh Tran, Giang Thi Hoang, Mathieu Dejean, Christelle Baptiste, Sovanna Tan, Laurent Laplaze, Jean Luc Verdeil, Pascal Gantet, Alexandre Grondin

## Abstract

Root anatomy plays a critical role in water and nutrients acquisition and influences plant adaptation to abiotic stresses. A better understanding of the genetic control of anatomical traits is important to improve rice production, a staple crop for half of the world’s population. In this study, we measured eleven root anatomical traits in a collection of 185 Vietnamese rice landraces, and conducted genome-wide association studies to uncover the genetic architecture underlying these traits. A total of 24 significant small nucleotide polymorphisms were identified, corresponding to 19 quantitative trait loci for traits such as root cross section area, cortex area, metaxylem area or theoretical axial conductance. Interestingly, several root anatomical traits correlated with previously characterized root architectural traits measured within the same Vietnamese rice collection, and some genetic regions associated with anatomical traits and drought-tolerance-related traits colocalized. Candidate genes potentially involved in root development and cell wall patterning were identified within these QTLs regions. In particular, the significant SNP in qRA10.1 associated with metaxylem area induces an amino-acid change in the kinase domain of a wall associated kinase (WAK) receptor-like protein kinase (OsWAK111) with an extracellular epidermal growth factor (EGT)-containing domain that bind pectin, which presumably sense perturbations in this component to influence cell elongation during plant development. This study reveal key determinants of the genetic control of root anatomy in rice, which could be exploited to enhance tolerance to abiotic stresses such as drought.

## Introduction

Plant roots are responsible for the absorption of water and nutrients from the soil. After absorption, solutes move radially from the epidermis to the xylem vessels, and then axially to the shoots. The radial flow of solutes has two possible paths, the apoplastic pathway through the cell walls and the symplastic pathway from cell-to-cell [1]. The axial flow depends on metaxylem vessel number and diameter [2]. Thus, tissue organization within the root can greatly impact the flow of water and nutrients to the shoots [3,4].

Rice (*Oryza sativa* L.) is a staple food for over half of the world’s global population, providing essential sustenance and nutrition to billions of people around the world. The rice root anatomy is typical of monocotyledonous plants, with an outer ground tissue formed of an epidermis, lignified exodermis and sclerenchyma cells, and several layers of cortex cells that can form aerenchyma [5]. The innermost tissue is composed of a stele, delimited by a suberized endodermis, and containing vascular tissues. As compared to other monocots, rice possesses a heavily lignified sclerenchyma layer between the cortex and the exodermis that restricts oxygen diffusion from the root to the soil, thereby limiting anoxic stress under flooded conditions [6]. Furthermore, aerenchyma are particularly developed in rice, also representing an adaptation to anoxic conditions by facilitating gas exchange with the shoot and providing a reservoir of oxygen for root cells respiration [7]. Aerenchyma, together with exodermis, sclerenchyma and endodermis have been proposed to limit the radial flow of water and nutrients from the outer ground tissue to the xylem [8,9]. However, rice is adapted to a wide range of agrosystems characterized by different types of soil and watering regimes. Therefore, rice root anatomy vary among different varieties, and these variations may be exploited to improve tolerance to abiotic stresses [10–12].

Drought is one of the major environmental factors limiting rice productivity and quality worldwide [13]. Several root anatomical traits have been linked to improved drought tolerance in cereal species [14]. In maize, an increase in root cortical aerenchyma and a decrease in cortical cell layers and size correlate with deeper root growth and greater water acquisition from drying soil [15–17]. The reduction of carbon investment into root tissue resulting from the presence of aerenchyma is considered beneficial for deeper root growth, as it supposedly allows for the reallocation of carbon to the growth of individual roots [18]. Similarly, an increase in cortical parenchyma cell wall width was associated with a reduced root metabolic cost and improved performance under drought in maize [19]. In addition, reduced root xylem vessel diameter was associated with yield increase in wheat under dry environments [20]. In rice, the relevance of these traits for drought tolerance remains to be clarified through field testing of appropriate genetic materials contrasting in these traits. This requires a comprehensive understanding of the genetic determinants controlling root anatomical traits. Although the gene network responsible for anatomical patterning has been well described in rice [21,22], the genes involved in the quantitative control of root tissue differentiation have been poorly described. Only few studies have explored rice root anatomical diversity at large scale to decipher the genetic determinants of anatomical traits through genome wide association studies (GWAS) [23–25].

In this study, we characterized the genetic diversity for root anatomical traits in a Vietnamese rice landraces collection [26]. This collection comprises 185 genotypes collected from diverse agrosystems across Vietnam, exhibiting substantial variation in morphological traits, including root thickness, which suggested corresponding diversity in root anatomical traits [27]. This genetic material represents a highly valuable resource for local breeding programs and may provide novel alleles, as only a small fraction of the Vietnamese rice genetic diversity has been exploited in previous association studies [28]. Crown roots from plants of this collection were sampled under field conditions in Vietnam. Anatomical traits were quantified on root histological cross sections, and association genetics on these traits were conducted using available genetic data. Several quantitative trait loci (QTLs) associated with root anatomical traits and potential causative candidate genes were then identified.

## Materials and Methods

### Plant materials

The rice collection used in this study covers 185 genotypes, comprising 182 Vietnamese landraces (114 indica, 62 japonica, and 6 admixed accessions) [26], along with 3 reference genotypes (Nipponbare, Azucena, and IR64). Seeds from these genotypes were officially provided by the Vietnam National Plant Genebank managed by the Plant Resources Center (Hanoi, Vietnam).

### Field experiment

Plants were grown in a rice field located at Hai Phong, Vietnam (coordinates: 20° 51’ 59’’ N, 106° 40’ 57’’ E) between July and November 2019. Seeds were sown in seedling beds for two weeks before being transplanted into an open field under lowland conditions. Genotypes were divided into groups based on their flowering duration: 67 genotypes were classified as early flowering (fewer than 85 days), 72 genotypes as medium flowering (85–119 days), 23 genotypes as late flowering (120–154 days), and 23 genotypes as very late flowering (more than 155 days). The flowering groups were divided into blocks, with 12 blocks for the early group, 12 for the medium group, 4 for the late group, and 4 for the very late group. Each genotype was planted in three replicated plots within its flowering group. Upon transplantation, a 75 cm spacing was applied between flowering groups and 50 cm between blocks within flowering groups. Within blocks, each plot covered 1 m² and contained 25 plants, spaced 25 cm apart both between and within rows. Four weeks after transplanting, five crown root segments of 3 cm length were collected near the stem base of one plant in each replicate. Roots were conserved in ethanol 50% (v/v).

### Root anatomical phenotyping

Out of the five sampled crown roots, three that appeared most similar in thickness and age were selected for each replicate. The root imaging protocol was adapted from [29]. Each root was divided into two parts and further cut by a scalpel blade into multiple segments. Root samples were transferred into a softening solution (89 % water:10 % glycerol:1 % tween) for a minimum of one day before being embedded in 5 % (w/v) agarose. After solidification, the agarose blocks were resized and glued on a vibratome plate (Thermo Scientific Microm). Root samples were sectioned into 100 μm-thick slices using an Hm650v vibratome (speed 30–50, frequency 40, amplitude 1.0). Individual slices were then collected with a fine brush and transferred to humidified slides with glycerol 50 % (v:v). Imaging of cell walls autofluorescence was performed using a NIKON Eclipse Ni-E epifluorescence microscope equipped with a ‘DAPI longpass’ filter cube (excitation filter: 330–380; suppression filter: 400; emission filter: 420-800). The objective used was a 10X Plan Apo λ 0.45 NA. In total, eighteen images (with a horizontal and vertical resolution of 0.8693 x 0.8693 µm²) of root transverse sections showing the highest quality in terms of tissue integrity and contrast were produced for each genotype (3 replicates, 3 crown roots per replicate and two images per crown root).

To quantify the anatomical traits, we used a modified version of PHIV-RootCell, a macro toolset within the Fiji software developed by [29]. Upon launching the program, the RGB image is converted into an 8-bit grayscale image, followed by the application of a Gaussian filter. The contrast is automatically enhanced, and the threshold can be manually adjusted at each step of the analysis to create the selection. This method was applied to calculate the surface area of different tissues (root, cortex, aerenchyma, stele and metaxylem). Diameters of sclerenchyma and endodermis cells were estimated by averaging the lengths of lines manually drawn through the centers of five different cells (including cell walls) within the respective layers. In total, eight anatomical traits were measured: root cross section surface area (Root_area) in mm^2^, cortex surface area (Cortex_area) in mm^2^, aerenchyma surface area (Aer_area) in mm^2^, stele surface area (Stele_area) in µm², diameter of sclerenchyma cell (Scl_D) in µm, diameter of endodermis cell (En_D) in µm, metaxylem surface area (Mx_area) in µm² and number of metaxylem (Mx_no) (S1 Fig). From these traits, we further derived the ratio of stele to cortex surface area (Stele_Cortex_ratio), the mean area of metaxylem (Mx_mean_area) in µm², and the theoretical root axial conductance (Kx). Kx was calculated using the modified Hagen-Poiseuille equation as in [30], where *ρ* is the fluid density (1000 kg m^-3^ for water at 20°C), *η* the viscosity of water (1 x 10^-9^ MPa s^-1^ at 20°C), and *d* the average metaxylem diameter that was derived from the mean area of metaxylem.

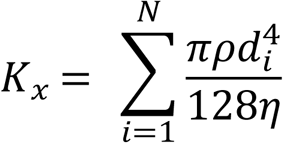

### Data analysis

All statistical analyses were performed using R Software (RStudio v.2023.03.0). The initial phenotypic dataset (raw dataset) contained a total of 18 images for each genotype. The raw dataset was cleaned out of outliers using analysis of variance (ANOVA; aov function) and Turkey HSD test (agricolae R package v.1.7.3) on each genotype for the root diameter trait. Values were considered outliers, and were removed for all other traits, if they differed significantly from the main set of values of root diameter measured for one genotype. For each trait, data from the different images were further averaged per repetitions, and a two-way ANOVA was performed to assess the effects of replication and genotype. Broad-sense heritability (*H*^2^) was determined according to [31] using the inti R package (v.0.6.8: Lozano-Isla, 2021) with the following formula:

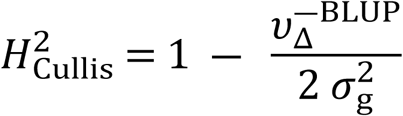

where *σ*^2^ refers to the variance, g to the genotype, 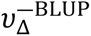 to the average standard error of the genotype best linear unbiased prediction (BLUP).

Trait mean value per genotype, standard deviations, and coefficients of variation for all traits were calculated. Mean values were used for Pearson’s correlation analysis (on the 185 genotypes using the “pairwise.complete.obs” option to account for NAs within the cor function), principal component analysis (on 132 genotypes after NAs removal) and association genetics analysis (on the 185 genotypes).

### Genome-wide association studies

Genome-wide association studies (GWAS) were conducted using the GAPIT3 package in R [32], using a previously published genotypic dataset consisting of 21,623 SNP (single nucleotide polymorphism) markers from 185 accessions within the Vietnamese rice collection [26]. The population structure had been previously analyzed, revealing six distinct genotypic groups [26]. Association analyses were performed using different complementary models: Mixed Linear Model (MLM), Generalized Linear Model (GLM), Multi-Locus Mixed-Model (MLMM), Conditional Masked Language Modeling (CMLM), and Bayesian-information and Linkage-disequilibrium Iteratively Nested Keyway (BLINK). SNPs were considered significant if their p-values were below the Bonferroni threshold calculated during the analysis [32]. Manhattan plots were generated using GAPIT3 [32].

### Linkage disequilibrium (LD) heatmap and identification of candidate genes

To define the QTL regions, pairwise linkage disequilibrium (LD) was calculated between significant SNPs and surrounding SNPs using the LDheatmap R package. The QTLs intervals were constrained to regions in which the squared allele frequency correlation (r²) between markers was greater than 0.4. If the initially defined region was smaller than 50 kb, it was extended to cover a 50 kb range both upstream and downstream of the significant markers. Genes within the QTLs regions were identified through the MSU Rice Genome Annotation Project Release 7 (http://rice.uga.edu/) and RAPDB version IRGSP-1.0 2023-09-07 (https://rapdb.dna.affrc.go.jp/). Candidate genes were then selected based on potential associations between their predicted function and the anatomical trait of interest. The expression levels of the selected candidate genes were obtained from RiceXPRo version 3.0 (https://ricexpro.dna.affrc.go.jp/index.html) [33].

## Results

### Variation in root anatomy in Vietnamese rice landraces

To analyse the diversity in anatomical traits in the Vietnamese rice collection, crown roots were sampled at the stem base of rice plants grown in the field in Vietnam. Root histological cross-sections were prepared, imaged, and analyzed using the semi-automated PHIV-RootCell developed under ImageJ. Trait variability ranged from 10% for the diameter of sclerenchyma cells to 86% for theoretical axial conductance, with traits related to root tissue area showing intermediate coefficients of variability (Table 1; S2 Fig). For instance, metaxylem area showed a phenotypic variance of 57.63% within the collection (Fig 1). Significant genotype effects (*p*-value < 0.001) were observed consistently across all traits, while repetition effects were limited to a few variables (diameter of endodermis cells and the mean area of metaxylem). Interestingly, high heritability (*H*² ≥ 0.82) was observed for all traits (Table 1). Overall, our data suggest that anatomical traits and theoretical axial conductance, that relate to root anatomical functions, were variable and highly heritable in the Vietnamese rice collection.

**Fig 1.**
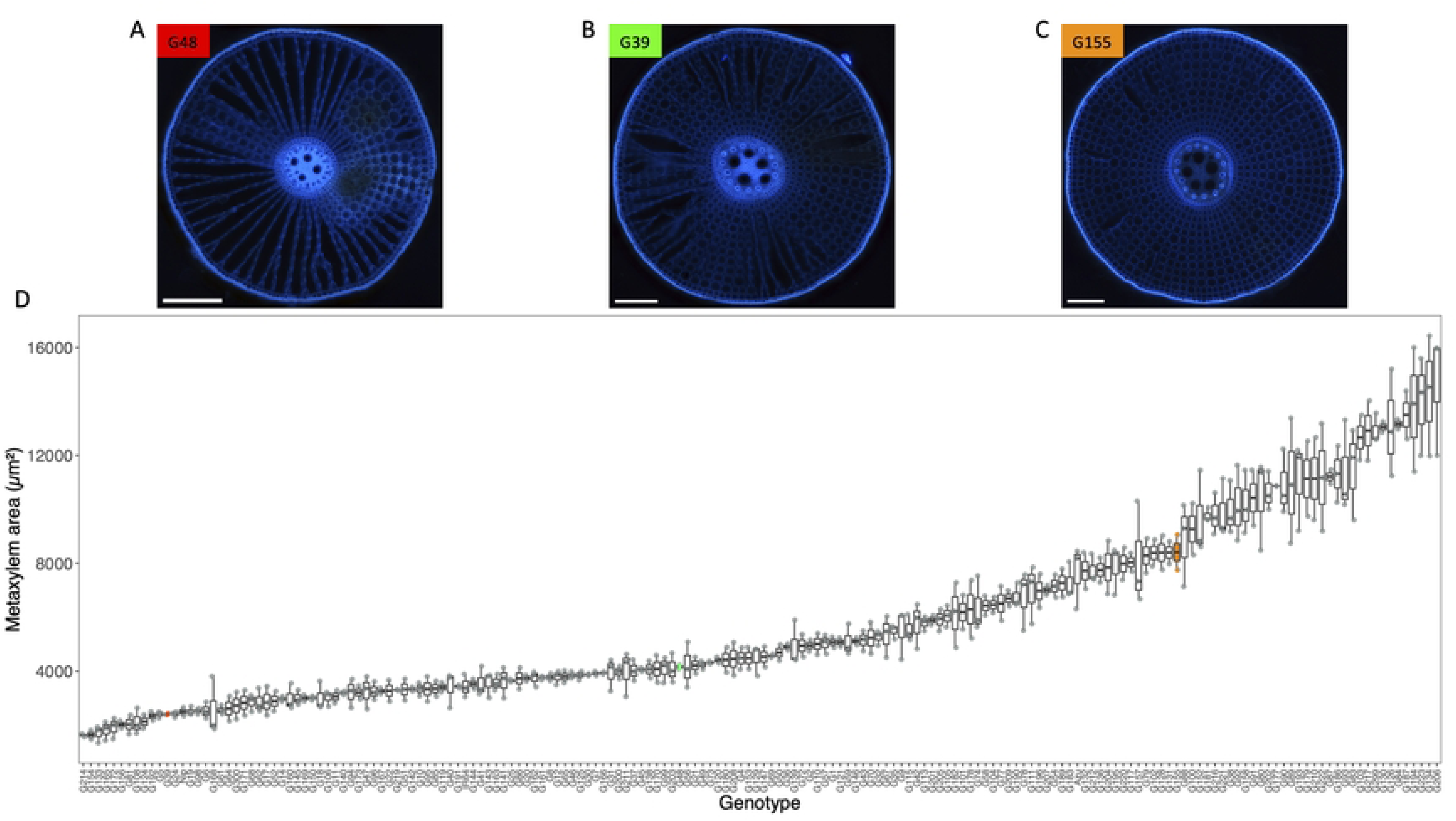
Variability for metaxylem area within the Vietnamese rice landraces collection. **A** to **C**. Cross section images of crown roots from three genotypes contrasting for root metaxylem area. For each genotype, the color boxes indicate the code of the different genotypes. **D**. Diversity for metaxylem area in the Vietnamese rice collection. The three genotypes represented in **A**-**C** are highlighted in E using the same color code.

**Table 1.** Phenotypic variation and broad sense heritability of root anatomical traits in the Vietnamese rice landraces collection.

| Traits | Mean | SD | Range |  | CV (%) | Replication | Genotype |  |
| --- | --- | --- | --- | --- | --- | --- | --- | --- |
|  |  |  | Min | Max |  | p-value | p-value | H<sup>2</sup> |
| Root_area (mm<sup>2</sup>) | 853451 | 276547 | 314618 | 1624432 | 32.40 | 0.382 | <0.001 | 0.92 |
| Cortex_area (mm<sup>2</sup>) | 766787 | 251887 | 277187 | 1453160 | 32.84 | 0.209 | <0.001 | 0.92 |
| Aer_area (mm<sup>2</sup>) | 322601 | 112489 | 109064 | 633661 | 34.86 | <0.05 | <0.001 | 0.83 |
| Stele_area (μm<sup>2</sup>) | 40757 | 17943 | 15277 | 104636 | 44.02 | 0.720 | <0.001 | 0.97 |
| Scl_D (μm) | 8.17 | 0.83 | 6.41 | 10.80 | 10.16 | 0.466 | <0.001 | 0.82 |
| En_D (μm) | 10.80 | 1.12 | 8.16 | 14.14 | 10.36 | <0.001 | <0.001 | 0.91 |
| Mx_area (μm<sup>2</sup>) | 5825 | 3356 | 1629 | 14645 | 57.62 | 0.111 | <0.001 | 0.97 |
| Mx_no | 4.57 | 0.92 | 2.61 | 7.66 | 20.18 | 0.507 | <0.001 | 0.94 |
| Mx_mean_area (μm<sup>2</sup>) | 1204 | 499 | 506.77 | 2395.65 | 41.44 | <0.05 | <0.001 | 0.95 |
| Stele_Cortex_ratio | 0.05 | 0.02 | 0.03 | 0.11 | 30.07 | 0.763 | <0.001 | 0.95 |
| Kx (kg m s<sup>-1</sup> MPa<sup>-1</sup>) | 6.75e-08 | 5.82e-08 | 1.02e-08 | 2.28e-07 | 86.33 | 0.064 | <0.001 | 0.93 |

SD: standard deviation; Min: minimum value observed in the collection; Max: maximum value observed in the collection; CV: coefficient of variation; H<sup>2</sup>: broad sense heritability. p-values from ANOVA tests are represented.

### Correlations between root anatomical traits, morphological traits and drought response-related traits

Pearson’s correlation analysis revealed several significant relationships among anatomical traits (Fig 2A). Most traits were significantly positively correlated with each other, except for the ratio between stele and cortex. The root cross-sectional area was highly correlated with the cortex area (*r* = 0.998) and less strongly correlated with the stele area (*r* = 0.72), indicating that cortex size is the primary determinant of overall root size. Furthermore, stele area showed strong positive correlation with metaxylem-related traits (surface area, mean surface area and number; *r* = 0.81 - 0.99). Theoretical axial conductance was almost perfectly correlated with mean metaxylem surface area (*r* = 0.99), highlighting the critical role of metaxylem size in water transport efficiency. By contrast, diameter of endodermis cells and of sclerenchyma cells had weak to moderate correlations with other traits, suggesting these traits may be regulated by distinct genetic and/or environmental factors.

**Fig 2.**
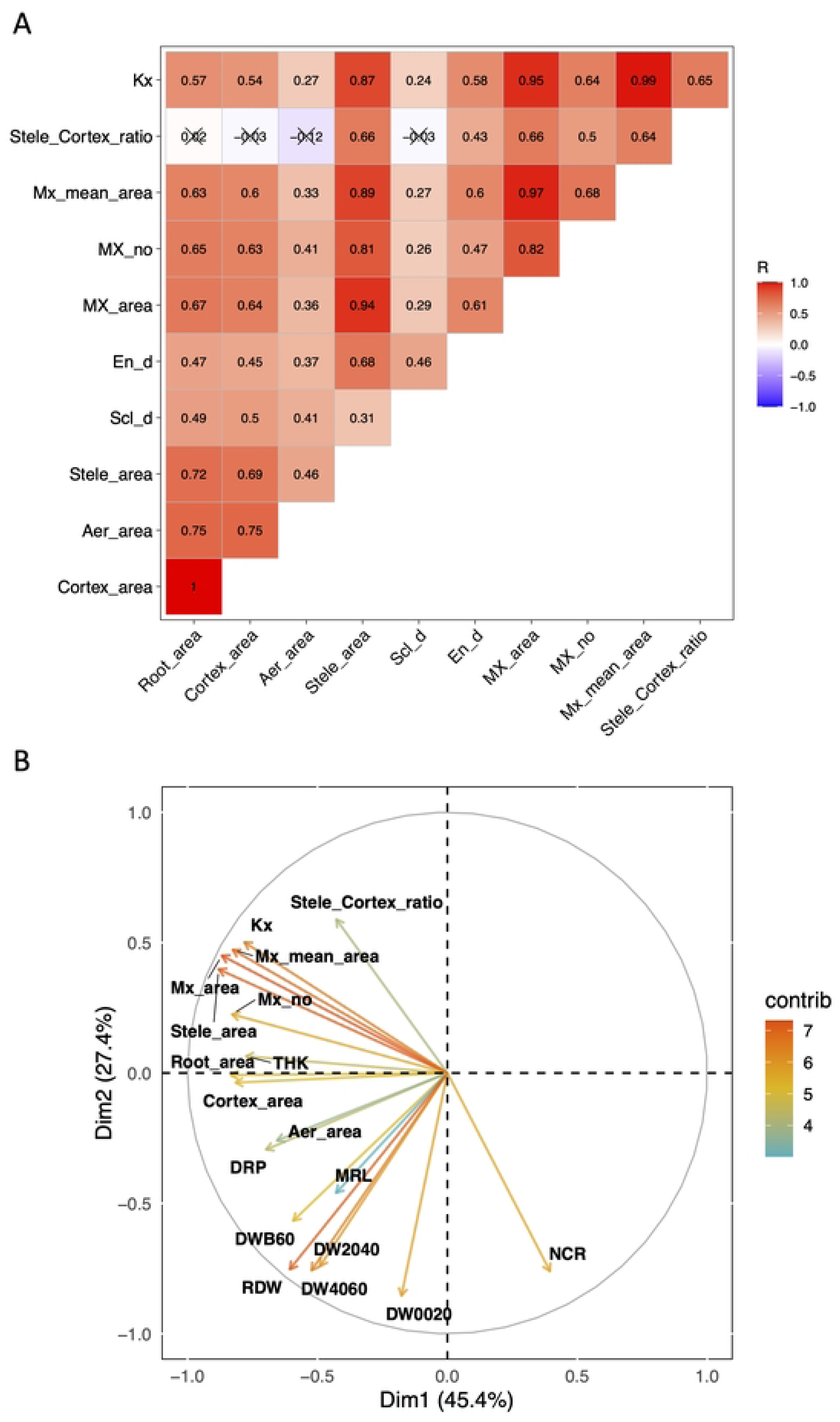
Correlation between root anatomical and architectural traits measured in the Vietnamese rice landraces collection. **A**. Pearson’s correlation coefficient (R) is indicated and non-significant correlations at a p-value threshold of 0.05 are indicated with a cross. **B**. Principal component analysis illustrating the relationship between anatomical and architectural traits. The contributions of the first two components are indicated and the color legend indicates the contribution of the different variables to the global variation. Root architectural trait data were retrieved from Phung et al., 2016. Mx: metaxylem; Kx: theoretical axial conductance; Aer: aerenchyma; THK: root thickness; DW0020: root mass in the 00-20 cm; DW2040: root mass in the 20-40 cm; DW4060: root mass in the 40-60 cm segment; DWB60: root mass below 60 cm; DRP: deep root proportion (< 40 cm); MRL: maximum root length; RDW: root dry weight; NCR: number of crown roots.

To obtain better insights into how anatomy and architecture integrate, we conducted correlation analyses between root anatomical traits measured in this study and 9 root architectural traits measured on plants growing in sand filled tubes by [27] using the same rice landraces collection. Most anatomical traits were positively correlated with root thickness (r = 0.44 to 0.67) but negatively correlated with crown root number (r = -0.27 to -0.62), except for aerenchyma area (S3 Fig). Root mass below 60 cm and deep root proportion (< 40 cm) showed consistent significant and positive correlations with anatomical traits (r = 0.21 to 0.46), except for the ratio of stele to cortex (S3 Fig). Conversely, the correlations were non-significant or negative between anatomical traits and root mass above 20 cm (r = 0.12 to -0.35), except for aerenchyma area (S3 Fig). Principal component analysis of root anatomical and architectural traits (Fig 2B) revealed distinct clustering patterns, with PC1 (45.4% variance) separating crown root number from all other traits (Fig 2B). The second component (explaining 27.4 % of the global variation) generally separated anatomical traits from architectural traits, except for cortex area and aerenchyma area that clustered with proportion of deep roots (Fig 2B). The intermediate positioning of theoretical axial conductance near anatomical traits aligns with its dependence on metaxylem vessel area, reinforcing its role as a proxy for hydraulic efficiency.

### Identification of QTLs related to root anatomy in Vietnamese rice landraces

To identify genomic regions associated with root anatomy, GWAS was conducted on the 11 anatomical traits using a previously published genotypic dataset composed of 21,623 SNPs [26] and different models [34]. Considering the Bonferroni threshold, a total of 46 associations comprising 24 SNPs were identified (S2 Table). Among these, 9 SNPs were identified across different GWAS models and 2 SNPs were identified across multiple traits (S2 Table).

Pairwise linkage disequilibrium was performed to delineate QTL around the significant SNPs. This analysis resulted in the identification of 19 QTLs scattered across 10 of the 12 rice chromosomes (Fig 3). Among the 19 identified QTLs, qRA6.2 on chromosome 6 associated with the stele area and contained five significant SNPs clustered together within a region of 234453 bp. Five QTLs contained at least two different SNPs associated with one or several traits and/or SNPs that were identified in at least two different GWAS models (qRA2.1, qRA6.1, qRA7.4, qRA8.1 and qRA10.1). SNPs present within these five robust QTLs explained relatively high phenotypic variance as compared to the other QTLs. Sj08_26825483R within qRA8.1 explained around 50% of the mean metaxylem area variance, with the major allele contributing to an increase of the trait of around 2000 µm² as compared to the minor allele (when averaged across all GWAS models). Similarly, Dj07_26030956R within qRA7.4 explained around 37% of the metaxylem area variance, with the major allele contributing to a reduction of the trait of around 2677 µm² (computed by MLMM).

**Fig 3.**
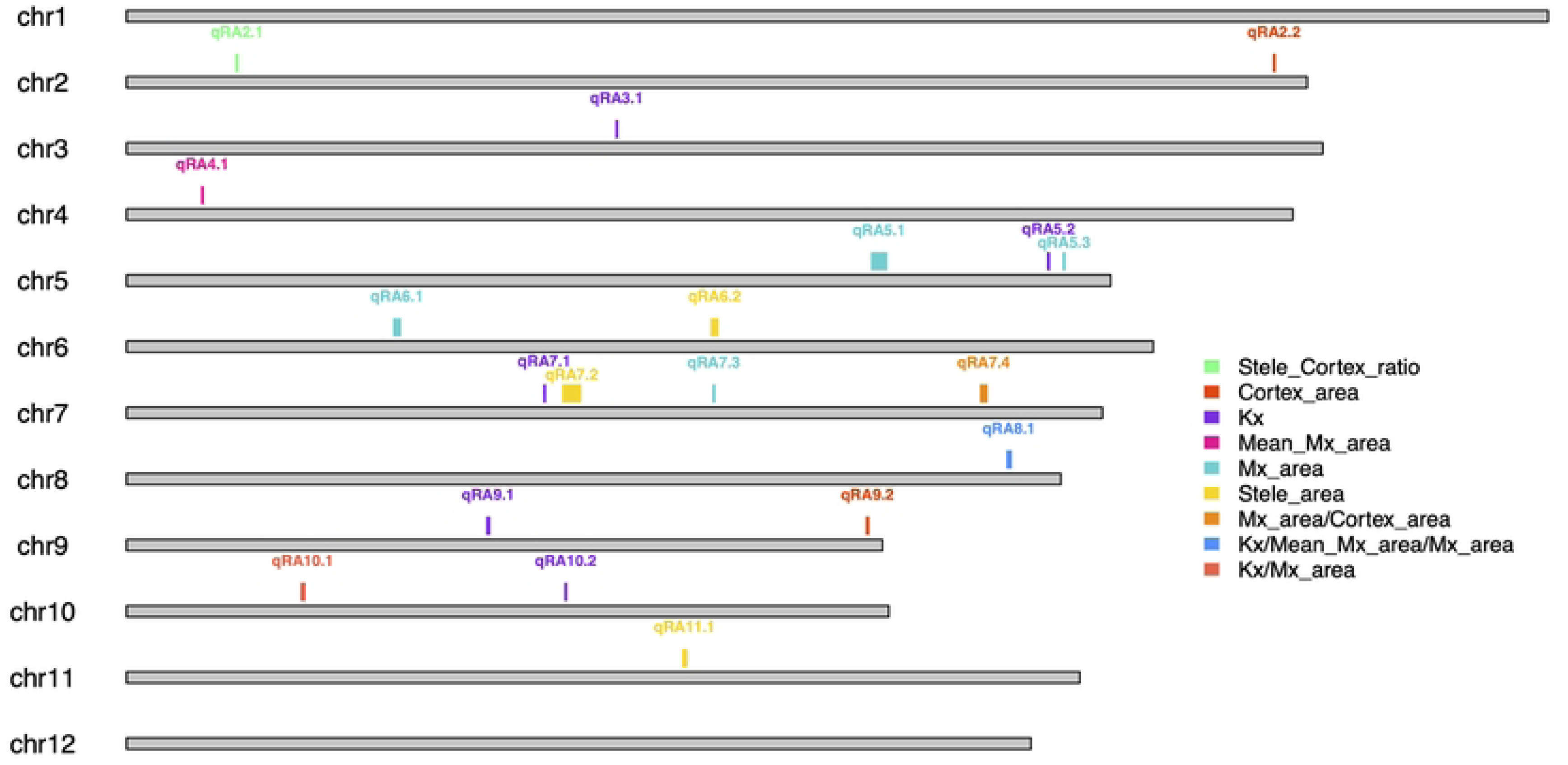
Karyotype representing the position of the 19 QTLs associated with root anatomical traits and theoretical axial conductance identified in the Vietnamese rice collection. The grey bar represents the chromosome length and the color code indicates the trait associated with the QTL.

We further examined whether QTLs for root anatomy identified in this study colocalized with QTLs previously identified for root anatomy in rice. qRA5.2 and qRA8.1 associated in our study with theoretical axial conductance and metaxylem area, colocalised with a QTL associated with stele diameter and diameter of metaxylem, respectively, identified in a panel of over 200 rice accessions from Southeast Asia [24]. However, none of the QTLs identified in this study colocalized with other QTLs associated with root anatomy identified in an indica panel [23], in the Rice Diversity Panel 1 [25] or in a biparental population created from a cross between IR64 and Kinandang Patong cultivars [35], indicating that the diversity in the Vietnamese collection allowed the identification of new QTLs controlling root anatomy.

We also examined whether root anatomy QTLs identified in this study colocalized with QTLs associated with plant and root architecture and traits associated with salt and drought stress tolerance that had been previously identified within the same collection of Vietnamese rice landraces [27,34,36–38]. qRA5.1, associated with metaxylem area, colocalized with a QTL associated with leaf water content under salinity stress [38]. Interestingly, leaf water content differed significantly between the two allelic group at the significant SNP within this QTL (S4 Fig). qRA6.2, associated with the stele area, colocalized with a QTL associated with plant growth under drought [37]. No differences in plant growth under drought was observed between the allelic groups at the significant SNPs within this QTL (S5 Fig). Last, qRA11.1, that is also associated with the stele area colocalized with a QTL associated with the number of tillers per plant [27]. Tiller number was significantly different between the two allelic groups at the significant SNP within this QTL (S6 Fig). Overall, these results suggest that our genetic analysis identified robust QTLs for rice root anatomy with strong linkage with the plant organ development and responses to abiotic stresses.

### Potential candidate genes related to root anatomy in Vietnamese rice landrace

To identify candidate genes associated with root anatomy in rice, we analyzed the genes present in the different QTLs regions. Our criteria for defining a gene as an interesting candidate were based on gene annotation, gene expression, gene distance to the most significant SNPs and known function in root development or root tissue differentiation in the literature. We highlight below five of the most interesting candidate genes that we identified (Table 2; S2 Table).

**Table 2.** List of candidate genes identified in QTLs associated with root anatomy traits.

| QTL | Trait | Chr | Gene ID | Gene annotation - function | Reference |
| --- | --- | --- | --- | --- | --- |
| qRA5.1 | Mx_area | 5 | LOC_Os05g39350/Os05g0470000 | OxXOAT2, xylanO-acetyltransferase, related to vasculature development | Zhong et al. 2018 |
| qRA7.2 | Stele_area | 7 | LOC_Os07g24190/Os07g0424400 | CESA3, cellulose synthase | Taylor et al. 1999; Lou et al. 2022 |
| qRA8.1 | Mx_area/Mx_mean_area/Kx | 8 | LOC_Os08g42470/Os08g0536800 | BEE1, BR Enhanced Expression 1, brassinosteroid signaling pathway | Wang et al. 2019; Saito et al. 2018 |
| qRA9.2 | Cortex_area | 9 | LOC_Os09g39220/Os09g0565600 | OsFackel1, delta14-sterol reductase, brassinosteroid synthesis | Jang et al. 2000; Li et al. 2024 |
| qRA10.1 | Kx/Mx_area | 10 | LOC_Os10g010030/Os10g0178800 | OsWAK111 - OsWAK receptor-like protein kinase, pectin sensing | Kanneganti & Gupta, 2011 |

Chr: Chromosome.

*LOC_0s05g39350* is located within qRA5.1 associated with surface area of metaxylem vessels, at 239925 bp downstream the significant SNP marker Dj05_22840944F explaining 19% of the phenotypic variance (S2 Table; S7 Fig) and encodes a xylan O-acetyltransferase 2 (XOAT2). This gene is strongly expressed in the roots during the vegetative growth phase, especially in the epidermis, exodermis and sclerenchyma in the elongation zone and early maturation zone [33]. In Arabidopsis, mutants defective in xylan acetylation exhibit a disorganised secondary wall structure, with collapsed xylem vessels and impaired plant development [39].

qRA7.2 is associated with the stele area and contains *LOC_Os07g24190* that encodes a protein belonging to the cellulose synthase family (S8 Fig). Within this QTL, the most significant SNP explaining around 30% of the phenotypic variance falls within an intergenic region at 16148 kb upstream of the start codon of *LOC_Os07g24190*. This gene is mostly expressed in roots and stems, more specifically within the stele of roots at the elongation and early maturation zone [33]. In *Arabidopsis thaliana*, mutation of a cellulose synthase gene, *CESA7*, causes defects in the xylem secondary cell wall formation and collapse of xylem elements [40,41], suggesting that cellulose synthase activity is linked to root anatomical development, with potential implications on the stele area.

In qRA8.1 that is associated with surface area and mean area of metaxylem vessels, as well as with theoretical axial conductance, we identified *LOC_Os08g42470*, encoding an orthologous to the basic/helix-loop-helix (bHLH) transcription factor BR ENHANCED EXPRESSION 1 (BEE1) protein from Arabidopsis (S9 Fig). The gene is located 15581 bp from the most significant SNP within this QTL (Sj08_26825483R), which explains on average 50% of the phenotypic variance (S2 Table). BEE1 is expressed in root tissues, especially in the differentiation and early maturation zones, although at significantly lower levels than in inflorescence [33]. Its expression is regulated by the BRI1-EMS-SUPPRESSOR 1 (BES1) transcription factor that functions as a key regulator in the brassinosteroid pathway in Arabidopsis [42]. BES1 also regulates root phloem and xylem development by promoting bidirectional differentiation of procambial cells into xylem and phloem cells in the root differentiation zone of Arabidopsis [43].

*LOC_Os09g39220*, also named *OsFACKEL1* and that encodes a delta14-sterol reductase involved in phytosterols and brassinosteroid biosynthesis [44], is an interesting candidate gene located within qRA9.2 associated with cortex area (S10 Fig). *OsFACKEL1* is located 60602 kb upstream of the most significant SNP explaining 20% of the phenotypic variance. This gene is most prominently expressed in roots, more particularly in the endodermis, pericycle, and stele regions of the elongation zone and early maturation zone [33]. Mutants of the orthologous gene in Arabidopsis, named *fackel*, present several defects in development, including abnormal cell division and expansion in cortical cell layers [45].

In qRA10.1 associated with metaxylem area and theoretical root axial conductance, we identified *LOC_Os10g010030* that encodes OsWAK111, a wall associated kinase (WAK) receptor-like protein kinase with an epidermal growth factor-like domain (Fig 4A-E). This gene is almost exclusively expressed in rice roots (Fig 4F). The most significant SNP that explains around 26% of the phenotypic variance falls within the last codon of the gene and causes an amino-acid change from Proline to Serine. WAK proteins play important functions in developmental processes, cell differentiation and stress responses [46]. Mutation in the WAK-like genes in tomato (*SlWAKL2*) or in Arabidopsis (*AtWAKL14*) induces a reduction in the number of xylem vessels compared to wild type plants [47].

**Fig 4.**
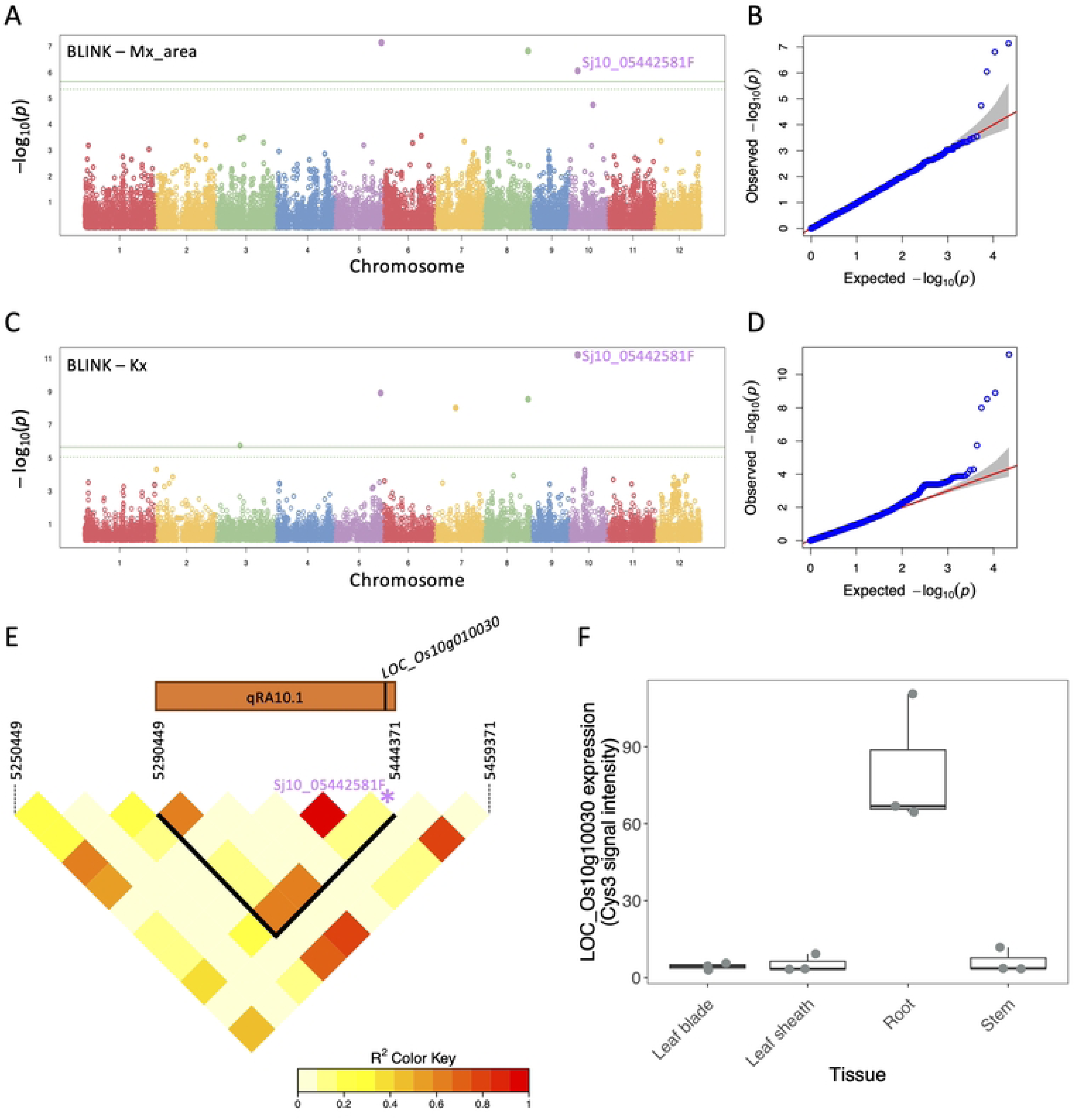
GWAS results for QTL qRA10.1 associated with metaxylem area and theoretical axial conductance (Kx). **A**, **C**. Manhattan plots generated from a GWAS performed using the BLINK model for metaxylem area and Kx. The solid green line represents the Bonferroni threshold and the green dashed line represents the suggestive significant threshold. **B**, **D**. Corresponding QQ-plots. **E**. Linkage disequilibrium plot for the QTL region spanning 153922 Kb, with an extension of 40 kb upstream and 15 kb downstream, with color shading representing the R^2^ values. The approximate position of the WAK111 gene (*LOC_Os10g10030*) within the QTL is marked and the significant SNP is highlighted in purple. **F**. Expression levels of *LOC_Os10g10030* in different tissues, retrieved from the RiceXpro database (Sato et al., 2013).

## Discussion

Root anatomy has a great influence on plant hydromineral nutrition. Thus, a better understanding of the genetic control of rice root anatomy will contribute to improve nutrient and water transport, with subsequent impact on tolerance to abiotic stresses such as drought. Despite its importance, the anatomical phenotypic landscape of rice and its genetic control remain poorly characterized, primarily because of the phenotyping bottleneck. In this study, we phenotyped a collection of 185 rice landraces originating from diverse geographic regions and ecosystems across Vietnam for root anatomical traits. Our findings confirm the diversity in root anatomical traits, provide insights on their interactions with plant and root architectural traits and drought and salinity tolerance, and uncover key genetic determinants of rice root anatomy, highlighting several interesting candidate genes.

Our results showed substantial variation in anatomical traits, supporting the large diversity within this Vietnamese rice collection previously observed for morphological [27,28,37], grain quality [34,48] and stress-related traits [37,38]. The range of variation in anatomical traits we measured was comparable to what has been previously observed by the few studies characterizing root anatomy at large scale within different rice diversity panels [23–25]. In order to express root anatomical traits in a more functional variable, we used the Hagen Poiseuille equation to compute the theoretical root axial conductance. The values we obtained were higher than those modelled for maize primary [49] and crown roots [50]. However, as the Hagen Poiseuille equation tends to overestimate axial conductance, cross-method comparisons of this parameter may be interpreted with caution. Most importantly, the root area measured in this study from field growing plants was positively associated with root thickness measured from plants growing in nethouses in sand filled tubes on the same rice collection by [27]. This suggests that root diameter is a robust trait that is relatively stable across growth environments. Overall, the variation in root anatomical traits measured in this study aligned with previous findings, and all traits showed high heritability, providing a strong foundation for association genetics analyses.

The Bonferroni threshold was used in our GWAS to select highly significant SNPs associated with root anatomical traits. This allowed us to delimit 19 QTLs associated with traits such as cortex area, stele area, metaxylem area, mean metaxylem area and theoretical root axial conductance. Using this threshold, no associations were found for root cross section area, aerenchyma area, number of metaxylem vessels and diameter of sclerenchyma cells and endodermis cells. Among the identified QTLs, two associated with metaxylem traits overlapped with QTLs previously reported for related traits in other rice diversity panels [24]. When compared with QTLs previously identified within the same Vietnamese landrace collection for other morphological and functional traits, we found that qRA6.2, associated with stele area, colocated with QTLs linked to growth under drought [37]. Stele area is strongly correlated with metaxylem area, a trait that modulates root water transport capacity, and that was associated with drought tolerance in several species [20,30,51]. Furthermore, most anatomical traits measured in this study were negatively correlated with shallow root growth and positively correlated with deep root growth measured in tubes [27]. Anatomical traits were also negatively correlated with the number of crown roots [27]. Overall, this suggests that, within the Vietnamese collection, genotypes with less crown roots of larger diameter, resulting from larger cortex area, stele area and metaxylem areas, may possess an enhanced capacity for deep root growth, conferring potential benefits for drought tolerance [11,52]. Further modeling and genetic studies are needed to confirm the interactions and synergies among these traits, investigate the potential additive effects of the underlying QTLs, and assess their impact on rice yield under varying drought conditions.

Brassinosteroids are important growth regulators, involved in many physiological processes such as cell expansion and division or vascular bundle differentiation [53]. Mutation of brassinosteroid receptors in Arabidopsis and rice showed that brassinosteroids are necessary for the normal development of xylem tissue [54,55]. In qRA8.1 associated with metaxylem size, we identified the BEE1, which function has been linked with metaxylem cell fate [43], as well as flavonoid accumulation under low temperature [56] and photoperiodic flowering in Arabidopsis [42]. To our knowledge, the BEE1 gene has not been functionally characterized in rice, although another gene called *Brassinosteroid Upregulated1* (*BU1*) encoding a bHLH protein phylogenetically close to BEE1, and involved in brassinosteroid signaling has been identified [57]. Overexpression of the *BU1* resulted in abnormalities in the pattern of internode elongation, tillers production and lignification and development of crown roots, but did not display morphological changes in vascular bundles [57]. Detailed analysis of root anatomical defects in BEE1 mutant will be necessary to confirm its role in metaxylem development. Similarly, we identified the *OsFACKEL1*, encoding a protein involved in the synthesis of brassinosteroids precursors [44], as a potential regulator of cortex area. A rice mutant for this gene has been characterized and exhibits abnormal shoot development, but no root phenotypes have been reported [44], underscoring the need for more in-depth investigations of root development defects.

Genes involved in cell wall patterning and sensing have also been identified close to associations related to stele surface area and metaxylem-related traits. In particular, wall-associated kinase (*OsWAK111*) was identified in qRA10.1 as a potential regulator of metaxylem area and theoretical root axial conductance (Kx). WAK are transmembrane proteins with possessing a signaling cytoplasmic Serine/Threonine kinase domain and an extracellular epidermal growth factor (EGT)-containing regions that bind pectin of multiple forms in the cell wall [46]. *WAK* genes presumably sense pectin perturbations to influence cell elongation during plant development [58–60]. Genes from this family are involved in xylem development in tomato and Arabidopsis (Ma et al., 2024). In rice, silencing of *WAK1* resulted in short and thicker roots without any root hairs [61], suggesting that it may impact root cell elongation, and potentially metaxylem area. The rice *OsWAK10* was identified as regulating sclerenchyma cell wall thickness and cell expansion and internodes length in stems, and allelic variations in the coding sequence of *OsWAK10* promoting shorter stem phenotypes seem to have been selected during domestication of japonica rice [62]. Interestingly, the significant SNP at qRA10.1 falls within the cytoplasmic kinase domain of OsWAK111 and induces an amino-acid change from Proline to Serine. We hypothesize that this change may alter the function of this protein, potentially disturbing signaling between the cell wall and cytoplasm with subsequent effect on cell elongation and metaxylem area. We also identified a xylan O-acetyltransferase 2 (XOAT2) in qRA5.1 associated with surface area of metaxylem vessels, and a cellulose synthase in qRA7.2 associated with the stele area. O-acetyltransferase proteins play important roles in O-acetylation of cell wall polymers, including xyloglucan, xylan, mannan, pectins and lignin [63], while cellulose synthases are required for cellulose biosynthesis [64]. These components are deposited in the cell wall and contribute to its structural strength and rigidity. Although mutants in these genes exhibit alterations in xylem morphology, their precise role in xylem development and function remain to be elucidated.

In conclusion, our study using a Vietnamese rice landrace collection identified several QTLs associated with root anatomical traits, some of them containing promising candidate genes which should be further investigated. Vietnamese landraces that possess QTLs showing significant variation in anatomical root traits, particularly in cortex area, stele and metaxylem area, could be valuable resources for improving rice abiotic and biotic stress tolerance, especially to drought and vascular pathogens.

## Acknowledgments

The Authors would like to thank the support of the University of Montpellier, the Institut de Recherche pour le Développement (IRD), the Centre de Coopération Internationale en Recherche Agronomique pour le Développement (CIRAD), and the Agricultural Genetics Institute of Hanoi. The authors would like to thank Nguyen Van Toan, Vu Thi Huong and Vu Manh An for their valuable support in field care and rice root harvesting. We acknowledge the Montpellier Ressource Imaging (MRI) facility, a member of the national infrastructure France-BioImaging for providing access to microscopy equipment and technical support.

## Author contributions

GTH performed the field experiment and root sampling. HLT performed the anatomical phenotyping, data analysis and genetic analysis under the supervision of MJ, CB, JLV, GTH, PG and AG. HLT, ST, LL, GTH, PG and AG contributed to the selection of candidate genes. GTH, PG and AG conceived the study and supervised the work. HLT, GTH, PG and AG wrote the manuscript. All authors read, edited and approved the final version of the manuscript.

## Data availability

All datasets supporting the finding of this study are available within the paper and its Supplementary Information.

## Abbreviation

qRA1.1: QTL identifier (q) for a root anatomy trait (qRA), located on chromosome 1 (qRA1), with the unique identifier number within this chromosome (qRA1.1).

## Supporting information

**S1 Fig. Root image analysis using PHIV-RootCell.**

**S2 Fig. Distribution of the different anatomical traits and theoretical axial conductance measured in the Vietnamese rice landraces collection.**

**S3 Fig. Correlation between anatomical and architectural traits measured in the Vietnamese rice landraces collection.** Pearson’s correlation coefficient (R) is indicated and non-significant correlations at a p-value threshold of 0.05 are indicated with a cross. Root architectural data were retrieved from Phung et al. (2016).

**S4 Fig. Repartion of surface area of metaxylem and leaf water content according to the allelic version at Dj05_22840944F in qRA5.1.** Surface area of metaxylem was measured in this study. Tiller number was retrieved in Hoang et al (2019). Stars indicate significance level. *: *p*-value < 0.05; ****: *p*-value < 0.0001.

**S5 Fig. Repartion of surface area of stele and relative crop growth rate (RCGR) according to the allelic version at Dj06_17819591R in qRA6.2.** Surface area of stele was measured in this study. Relative crop growth rate was retrieved from Hoang et al (2019). The repartion of relative crop growth rate was also studied in the other significant SNPs in this QTL (Dj06_17906389R; Sj06_17951687F; Sj06_17959313F; Sj06_17969293F). **: *p*-value < 0.01; ns: non-significant.

**S6 Fig. Repartion of surface area of stele and tiller number according to the allelic version at Dj11_16929520R in qRA11.1.** Surface area of stele was measured in this study. Tiller number was retrieved from Phung et al (2016). ****: *p*-value < 0.0001.

**S7 Fig. GWAS results for QTL qRA5.1 associated with metaxylem surface area. A**. Manhattan plot generated from a GWAS performed using the MLMM model formetaxylem surface area. The solid green line represents the Bonferroni threshold and the green dashed line represents the suggestive significant threshold. **B**. Corresponding QQ-plot. **C**. Linkage desequilibrium plot for the QTL region spanning 504443 Kb, with color shading representing the R^2^ values. The approximate position of the xylanO-acetyltransferase gene (*LOC_Os05g39350*) within the QTL is marked and the significant SNP is highlighted in blue. **D**. Expression levels of *LOC_Os05g39350* in different tissues, retrieved from the RiceXpro database (Sato et al., 2013).

**S8 Fig. GWAS results for QTL qRA7.2 associated with stele area. A**. Manhattan plot generated from a GWAS performed using the MLMM model for stele area. The solid green line represents the Bonferroni threshold and the green dashed line represents the suggestive significant threshold. **B**. Corresponding QQ-plot. **C**. Linkage desequilibrium plot for the QTL region spanning 582971 Kb, with color shading representing the R^2^ values. The approximate position of the cellulose synthase gene (*LOC_Os07g24190*) within the QTL is marked and the significant SNP is highlighted in blue. **D**. Expression levels of *LOC_Os07g24190* in different tissues, retrieved from the RiceXpro database (Sato et al., 2013).

**S9 Fig. GWAS results for QTL qRA8.1 associated with surface area, mean area of metaxylem vessels and theoretical axial conductance. A**. Manhattan plot generated from a GWAS performed using the BLINK model for surface area of metaxylem. The solid green line represents the Bonferroni threshold and the green dashed line represents the suggestive significant threshold. **B**. Corresponding QQ-plot. **C**. Linkage desequilibrium plot for the QTL region spanning 174319 Kb, with color shading representing the R^2^ values. The approximate position of the BR Enhanced Expression 1, gene (*LOC_Os08g42470*) within the QTL is marked and the significant SNP is highlighted in blue. **D**. Expression levels of *LOC_Os08g42470* in different tissues, retrieved from the RiceXpro database (Sato et al., 2013).

**S10 Fig. GWAS results for QTL qRA9.2 associated with cortex area. A**. Manhattan plot generated from a GWAS performed using the BLINK model for cortex area. The solid green line represents the Bonferroni threshold and the green dashed line represents the suggestive significant threshold. **B**. Corresponding QQ-plot. **C**. Linkage desequilibrium plot for the QTL region spanning 119793 Kb, with color shading representing the R^2^ values. The approximate position of the OsFackel1 gene (*LOC_Os09g39220*) within the QTL is marked and the significant SNP is highlighted in blue. **D**. Expression levels of *LOC_Os09g39220* in different tissues, retrieved from the RiceXpro database (Sato et al., 2013).

## Notes

### Competing Interest Statement

The authors have declared no competing interest.

